# Whale watching with BulkVis: A graphical viewer for Oxford Nanopore bulk fast5 files

**DOI:** 10.1101/312256

**Authors:** Alexander Payne, Nadine Holmes, Vardhman Rakyan, Matthew Loose

## Abstract

**Motivation:** The Oxford Nanopore Technologies (ONT) MinION is used for sequencing a wide variety of sample types with diverse methods of sample extraction. Nanopore sequencers output fast5 files containing signal data subsequently base called to fastq format. Optionally, ONT devices can collect data from all sequencing channels simultaneously in a bulk fast5 file enabling inspection of signal in any channel at any point. We sought to visualise this signal to inspect challenging or difficult to sequence samples, or where flow cell performance is modified by an external agent, such as ‘Read Until’.

**Results:** The BulkVis tool can load a bulk fast5 file and overlays MinKNOW classifications on the signal trace. Users can navigate to a channel and time or, given a fastq header from a read, jump to its specific position. BulkVis can export regions as Nanopore base caller compatible reads. Using BulkVis, we find long reads can be incorrectly divided by MinKNOW resulting in single DNA molecules being split into two or more reads. The longest seen to date is 2,272,580 bases in length and reported in eleven consecutive reads. We provide helper scripts that identify and reconstruct split reads given a sequencing summary file and alignment to a reference. We note that incorrect read splitting appears to vary according to input sample type and is more common in ‘ultra long’ read preparations.

**Availability:** The software is available freely under an MIT license at https://github.com/LooseLab/bulkVis. The software requires python3 to run.

## Introduction

Oxford Nanopore Technologies (ONT) range of sequencing platforms (MinION, GridION, and PromethION) utilise biological nanopores, embedded in a synthetic membrane, to sequence individual single stranded molecules of DNA ^1^. A potential is applied across the membrane and creates a current flow through the nanopore. As single stranded DNA passes through the aperture of the nanopore it creates characteristic disruptions in current flow dependent on the specific sequence in the pore at that moment ^2^. The real time nature of nanopore sequencing means that reads are written to disk as soon as the DNA has translocated through the pore. This uniquely enables rapid analysis of sequence data ideal for both field and clinical work ^3,4^. To do this, the software controlling sequencing (MinKNOW) monitors the state of each channel in real time to determine if the signal observed represents nucleic acid. MinKNOW processes the continuous data stream from the MinION device into individual read fast5 files that contain the raw signal data. These files are subsequently base called to retrieve the underlying sequence. At its most real time, the sequence of the DNA can be determined as the molecule is passing through the pore, enabling approaches such as ‘Read Until’ where specific molecules can be dynamically rejected from the sequencer according to user customisable parameters ^5^.

The real time partitioning of the data stream into reads results in the loss of information about the current state both before and after a read as these events are not recorded in a read fast5 file. To better understand these events and to view the effects of user intervention on sequencing when developing methods for read until or using difficult samples, we wished to visualise the entire data stream from the MinION device. ONT provide an optional bulk fast5 file format to capture the full complement of data from every channel on the sequencing device ^6^. The bulk fast5 file includes raw signals for every channel and metadata including the classifications made by MinKNOW on the raw signal stream (see Supplementary Table 1). To visualise bulk fast5 files, we developed BulkVis using the bokeh visualisation package ^7^. BulkVis can annotate signal features based on the metadata within the bulk fast5 file and provides a simple method to relate a base called read back to the channel and time in the data stream from which it originated. BulkVis also provides a feature to create read fast5 files from a selected region of a bulk fast5 file. These reads can then be called by a Nanopore compatible base caller.

In the course of developing BulkVis, we observed examples of reads incorrectly segmented by MinKNOW leading to a reduction in the read lengths reported. This incorrect splitting of reads appears to correlate with read lengths such that ultra long reads (colloquially referred to as ‘whales’) are more likely to be affected. In some cases there is no apparent reason for the read to have been split, but in many others we observe examples of reads that exhibit unusual signal patterns prior to the incorrect split.

## Results

### BulkVis

BulkVis runs as a bokeh app and scans a folder containing bulk fast5 files at startup. Individual files can be selected through the interface and specific channels plotted to the screen (Figure 1). Basic metadata associated with the bulk fast5 file are displayed to the user. To navigate the bulk fast5 file a specific region can be input as channel and time coordinates, in the format `channel: start time-end time`. Alternatively copying and pasting the fastq read header from a base called read will cause BulkVis to display the specific channel and time, for this read, in the bulk fast5 file. These files can also be navigated by jumping to the next or previous instance of a specific annotation, such as ‘strand’, meaning the nanopore is actively sequencing, or ‘pore’ meaning that the pore is open and available to sequence another molecule (see Supplementary Table 1). These annotations are overlaid on the signal plot as vertical dashed lines and annotated with the type and associated ID if available (Figure 1). The raw signal data are smoothed depending on the length of signal data being displayed to aid rapid visualisation. BulkVis also allows the user to export the signal section of the current position to a read fast5 file compatible with Nanopore base callers. To avoid confusion with MinKNOW derived reads, BulkVis reads are named based on the channel from which they are derived coupled with the start and end index of the read segment recorded in samples. As an example, the read segment shown in Figure 1 results in a read fast5 file named `plsp57501_20170308_fnfaf14035_mn16458_sequencing_run_nott_hum_whir_s2_60428_bulkvis-read_22448000-25724000_ch_450.fast5`. This region captures three single reads. When called as a single read, this region generates a sequence of 215,662 bases in length (Supplementary File Collection 1). The original three reads base call at a combined length of 215,153 bases in length (Supplementary File Collection 1) and the single called read maps well to the combined original three reads (Supplementary Figure 1).

**Figure 1.**
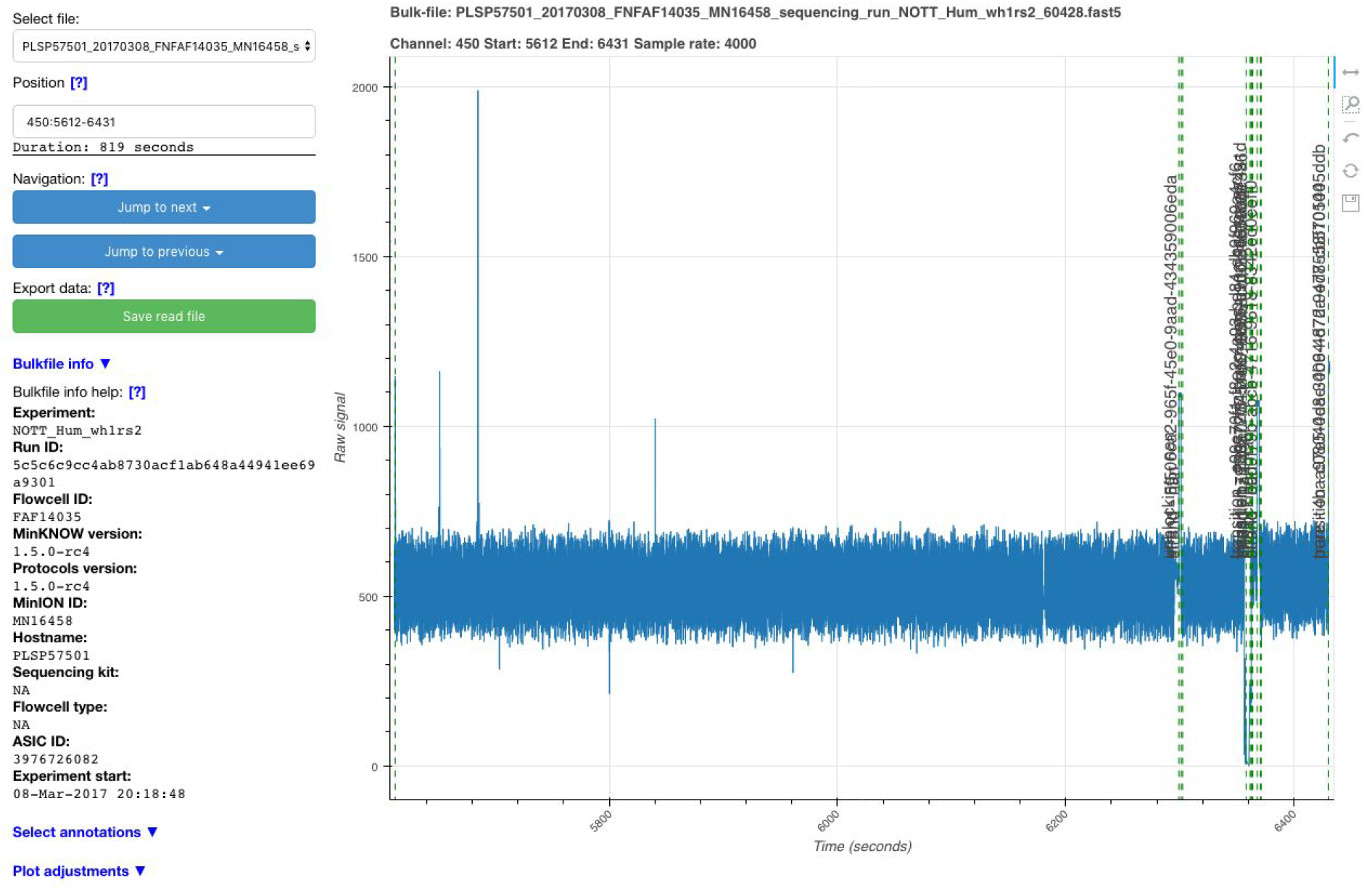
Screenshot of the BulkVis application running. The vertical dashed lines indicate different annotations overlaid by MinKNOW on the signal trace in real time. The left panel provides configuration and navigation options for the selected bulk fast5 file.

During MinION library preparation, adapter sequences are added to DNA molecules and so every sequenced read should begin with an adapter sequence. MinKNOW recognises these sequences in real time and usually labels a read start with the annotation ‘adapter’. A channel without DNA in a pore will have a current trace labelled ‘pore’. Then an adapter sequence should be detected (labelled ‘adapter’) followed by the signal derived from the read itself (‘strand’) (Figure 2A). BulkVis was developed in part to observe the effects of unblocking, the reversal of voltage across a specific channel to eject material from the pore, on DNA sequence as it traverses through a nanopore. Unblocking is used in two ways on nanopore sequencers; firstly the sequencer detects and removes blockages in the pore and, secondly, for the rejection of unwanted DNA in selective sequencing or ‘Read Until’ 5. The only way to observe the effect of an unblock on a channel immediately after the read has been ejected is to analyse a bulk fast5 file or to inspect the reads in order from an individual channel. An example unblock is shown in Figure 2B. At the time of writing, unblocks appear to have a fixed duration of ^2^ seconds after which the channel should return to its normal state. ONT are soon to release an updated version of unblock, termed “Progressive Unblock” which will gradually increase the duration of the flick time (MinKNOW 2.0 Stuart Reid Pers Comm.).

As part of our recent efforts to sequence the human genome on a MinION device ^8^, we generated a protocol to sequence ultra-long DNA molecules ^9^ and so we used BulkVis to investigate the signal from MinKNOW during one of these runs (ASIC ID 3976726082, Supplementary Note 1). We were surprised to observe a number of reads that did not show the expected ‘pore’, ‘adapter’, ‘strand’ sequence. We found ‘strand’ sequences that were separated by either an ‘above’ and/or ‘transition’ (Figure 2C) or even ‘unblock’ (Figure 2D) signals without any evidence of ‘pore’ or ‘adapter’ sequences present. We were surprised to see these events, reasoning that every sequenced read should begin with an adapter. We therefore closely examined the reads before and after these unusual read split events. For example, by looking at the mapping of the reads prior and post the events shown in Figures 2C and D, we determined that in both cases the two sequences were derived from adjacent positions on the same chromosome (Table 1). These reads, sequenced one after another, must presumably be derived from a single molecule. The alternative explanation is the chance sequencing of two independent molecules, one after another, through the same pore that map adjacently on the human reference.

**Figure 2.**
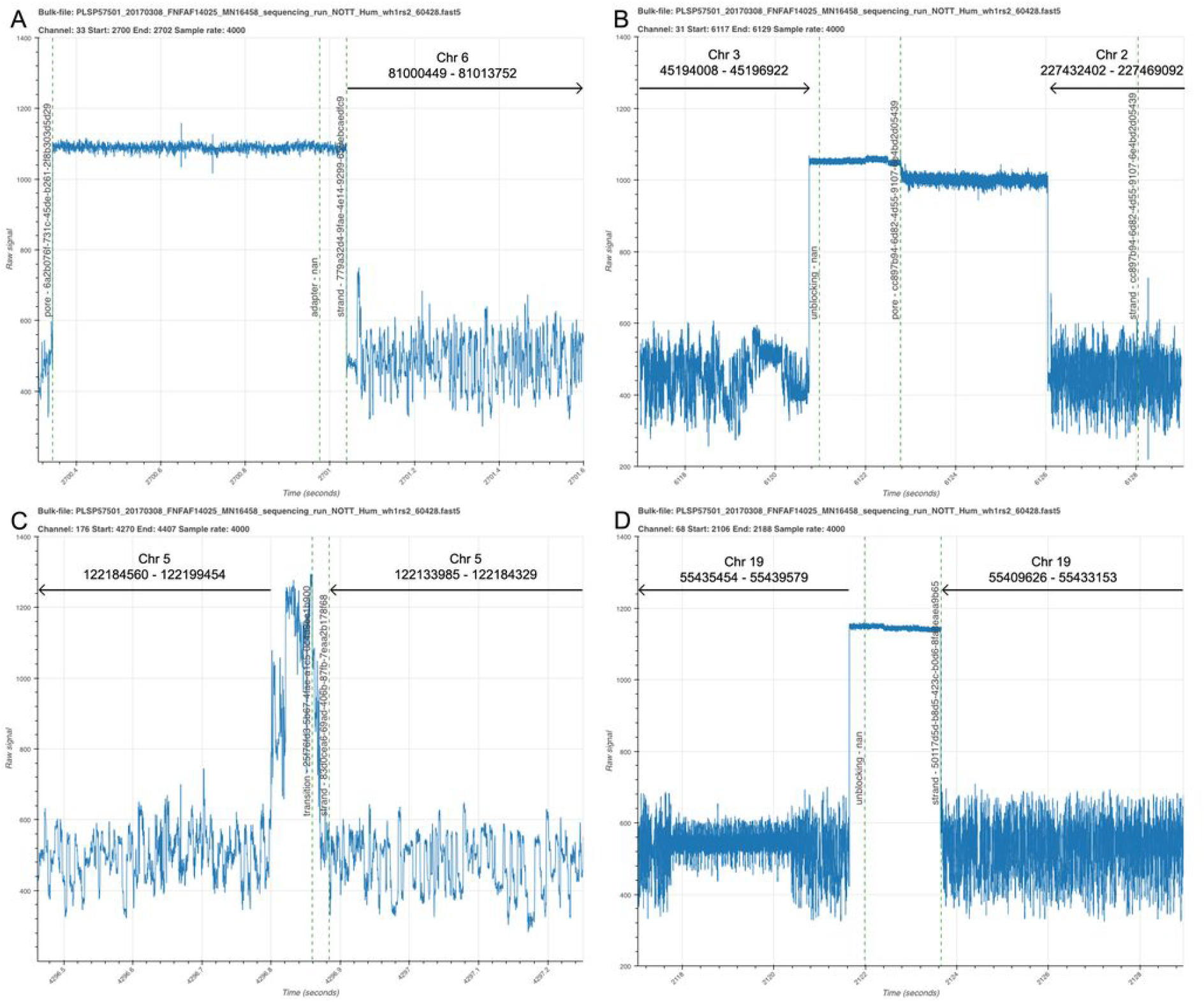
Illustrative segments from a bulk fast5 file visualised with BulkVis. (A) The start portion of a read mapping to chromosome ^6^. The open channel ‘pore’ state is shown followed by the detection of an ‘adapter’ sequence by MinKNOW and then the ‘strand’ annotation representing sequence signal. (B) A read which ends with an ‘unblock’ followed by open channel ‘pore’ and then a new sequencing read. (C) Two adjacent reads from the same channel separated by an unusual current pattern. Although these two reads are reported as distinct molecules by MinKNOW, they map consecutively to the reference. (D) Two adjacent reads separated by an ‘unblock’ signal. In this case, the unblock does not successfully remove the DNA. Instead the read continues to sequence again mapping adjacently to the reference.

**Table 1:** Mapping data for events show in Figure 2C,D from ASIC ID 3976726082. Reads mapped to GRCh38 using minimap2 -x map-ont. The combined read length for 2C is 56,284 bases and maps to a total span of 65,469 bases. The combined read length for 2D is 30,664 bases and maps to a total span of 29,953 bases. All reads shown here map in the reverse orientation.

|  | Read ID | Chan | Read | Length | Chr | Start | End |
| --- | --- | --- | --- | --- | --- | --- | --- |
| 2C | 7ed4aafb-d058-481c-ad60-903fd8327240 | 176 | 943 | 10,275 | 5 | 122184560 | 122199454 |
|  | 83d0cea6-69ad-406b-87fb-7eaa2b178f68 |  |  | 43,145 |  | 122133985 | 122184329 |
| 2D | c13c1e73-f7e0-4ae2-8cda-729f3b4dcb79 | 68 | 758 | 5,068 | 19 | 55435454 | 55439579 |
|  | 50117d5d-b8d5-423c-b0d6-8fa8eaea9b65 |  |  | 25,596 |  | 55409626 | 55433153 |

Given the unlikely nature of this event, we asked how many other reads in the same sequencing run showed this phenomenon. To do this we mapped all the reads from this single run (ASIC ID 3976726082) against the GRCh38 reference genome ^10^. We then used the read and channel numbers to sort the reads according to the order they had passed through each channel. We asked if adjacent reads mapped to contiguous positions on the reference genome (using the script whale_watch.py). From 75,689 total reads, 2,982 were incorrectly split with pairs of reads mapping adjacently to the reference. Simply stitching these reads together and recalculating the read length N50 resulted in an increase from 98,876 to 103,925 bases. The mean read length of the incorrectly split reads (55,190 bases) is higher than that of the entire dataset (23,717 bases). Re-examining all our previous ultra-long datasets revealed incorrect read splitting occurred from 1-10% of the time (Table 2). From any given run incorrectly split reads had consistently higher mean read lengths than those which appear to be true single molecules. As such, these reads have a significant effect on read N50, increasing our N50 measurements from previous runs by as much as 21 kb. We provide an accompanying script, whale_merge.py, to join putative split reads based on mapping to a suitable reference genome into renamed fastq files for downstream analysis.

**Table 2:** Read length statistics for ^14^ runs from Jain et al^8^z with incorrectly split reads calculated using whale_watch.py after mapping to GRCh38 using minimap2 -x map-ont.

| Asic ID | Read count |  |  | Mean |  |  | N50 |  |  |
| --- | --- | --- | --- | --- | --- | --- | --- | --- | --- |
|  | Original | Split | % | Original | Split | Corrected | Original | Corrected | Increase |
| 16056159 | 82,136 | 3,953 | 4.81 | 22,532 | 64,810 | 23,134 | 126,793 | 138,627 | 11,834 |
| 17958431 | 53,720 | 1,539 | 2.86 | 24,431 | 41,913 | 24,804 | 84,015 | 85,947 | 1,932 |
| 2901545329 | 41,384 | 932 | 2.25 | 20,299 | 51,910 | 20,534 | 59,500 | 61,168 | 1,668 |
| 3439856925 | 19,673 | 908 | 4.62 | 31,962 | 37,958 | 32,738 | 132,277 | 135,990 | 3,713 |
| 3709819546 | 73,752 | 2,489 | 3.37 | 28,268 | 56,948 | 28,777 | 129,792 | 135,156 | 5,364 |
| 3976726082 | 75,689 | 2,982 | 3.94 | 24,957 | 55,190 | 25,482 | 98,876 | 103,925 | 5,049 |
| 4109802543 | 61,223 | 2,769 | 4.52 | 26,129 | 59,149 | 26,776 | 114,934 | 123,304 | 8,370 |
| 4111860526 | 65,138 | 4,193 | 6.44 | 26,340 | 49,005 | 27,271 | 102,785 | 111,444 | 8,659 |
| 4178920553 | 270,189 | 12,045 | 4.46 | 10,680 | 14,967 | 10,936 | 26,744 | 27,759 | 1,015 |
| 4244782843 | 9,663 | 882 | 9.13 | 35,380 | 63,434 | 37,242 | 110,455 | 125,144 | 14,689 |
| 4245291640 | 72,931 | 6,860 | 9.41 | 21,243 | 55,293 | 22,410 | 102,621 | 123,768 | 21,147 |
| 4249180049 | 68,167 | 1,209 | 1.77 | 26,477 | 71,002 | 26,722 | 132,550 | 136,916 | 4,366 |
| 82266371 | 71,150 | 2,687 | 3.78 | 25,611 | 54,145 | 26,152 | 129,656 | 137,644 | 7,988 |
| 87644245 | 451,019 | 2,697 | 0.6 | 8,475 | 10,554 | 8,501 | 14,957 | 15,016 | 59 |

We have recently generated additional ultra-long reads derived from the same reference human genomic DNA sample using the updated RAD004 transposase kit for ultra long reads ^8,11^. We ran this same analysis on these reads and found the occurrence of incorrectly split reads to be higher than our original ultra-long read set, with up to 30% of reads in one run being affected and increases in read N50 of up to 40kb (data not shown). There are many differences between these runs including the input DNA, the sequencing kit and other unknown variables within the flowcells and MinKNOW software itself. Within this dataset we found a single read of 1,204,840 bases that maps to 1,325,742 bases on chromosome 5 (Figure 3A). More remarkably, we found a set of eleven reads which, when merged, were 2,272,580 bases in length. This merged read maps to a single location in the human genome spanning 2,290,436 bases (see Table 3, Figure 3B, Supplementary File Collection 2). Unfortunately, we did not collect a bulk fast5 file for this run. However, the next longest ‘fused’ read caught in a bulk fast5 file was 1,385,925 bases in length, derived from nine individual reads (see Table 4, Figure 3C, Supplementary Figure 2). Using BulkVis we created a single read fast5 file from the signal covering all these reads and base called it using albacore. This resulted in a similar read length and a read which maps in its entirety to a single location in the genome.

**Figure 3.**
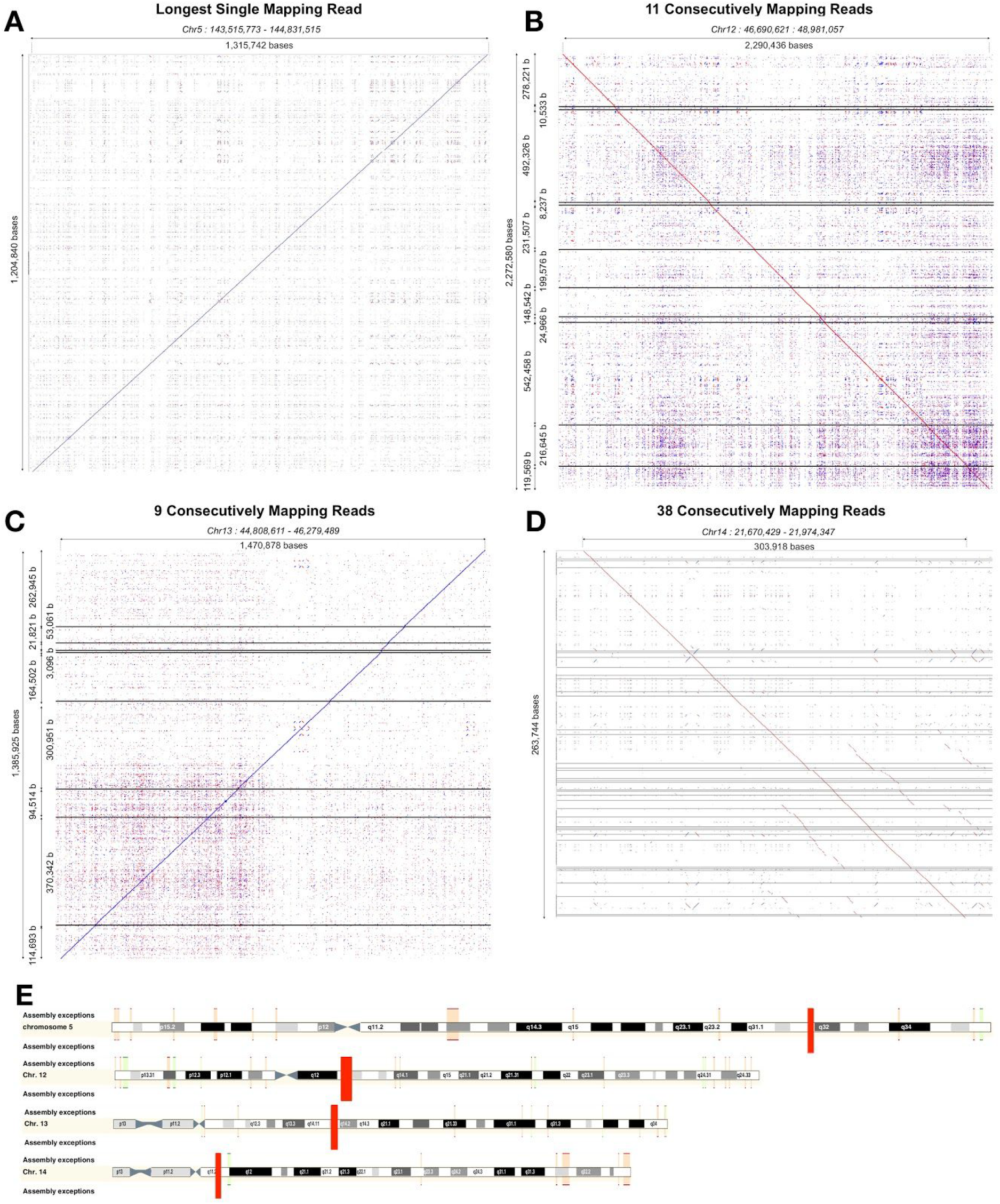
Read mappings. (A) The longest single read observed by us to date. (B) The longest fused read to be sequenced exceeding 2 Mb in length, caught in 11 reads. (C) The longest fused read sequenced for which we have the underlying bulk fast5 file. (D) A fused read comprising 38 individual sequences. (E) Illustration of the mapping of reads from A-D against the reference (GRCh38). Reads mapped and visualised with last (-m 1) and last-dotplot12. Horizontal lines indicate breaks between individual reads.

**Table 3:** Mapping data for eleven consecutive reads. Reads mapped to GRCh38 using minimap2 -x map-ont. The combined read length is 2,272,580 bases and maps to a total span of 2,290,436 bases.

| Read ID | No | Ch | Len | Chr | Std | Start | End |
| --- | --- | --- | --- | --- | --- | --- | --- |
| cbb741c3-27da-477e-87c8-20d91016251d | 187 | 396 | 278,221 | 12 | + | 46690621 | 46969898 |
| 7d6f6ae9-68c1-4b1e-84b5-a7aab0758c5b | 191 | 396 | 10,533 | 12 | + | 46970132 | 46979349 |
| f10eac6e-03ec-43fe-848a-769c3c85edcc | 195 | 396 | 492,326 | 12 | + | 46980918 | 47477341 |
| a19c94a3-4798-4422-a770-3122ce343de7 | 205 | 396 | 8,237 | 12 | + | 47477770 | 47485893 |
| 7c39fe08-b7d5-4160-831d-4364c7871fc4 | 206 | 396 | 231,507 | 12 | + | 47486059 | 47719483 |
| 109c8239-7357-4c20-b3a6-c1b7a95e0b48 | 208 | 396 | 199,576 | 12 | + | 47719572 | 47920652 |
| 0e63d0f4-0e41-4eb8-bfd2-69143efb70f5 | 211 | 396 | 148,542 | 12 | + | 47920996 | 48071262 |
| 760d1f30-ac17-4026-84e5-a94856c6d650 | 212 | 396 | 24,966 | 12 | + | 48071616 | 48096201 |
| ac191a38-ce73-4610-ae5b-8fbe03178ad7 | 216 | 396 | 542,458 | 12 | + | 48096764 | 48640714 |
| 07df9239-2302-436c-9430-048c6061c848 | 223 | 396 | 216,645 | 12 | + | 48640869 | 48859917 |
| c371771b-016c-4bfc-9dae-592e6631d0ba | 231 | 396 | 119,569 | 12 | + | 48860239 | 48981057 |

**Table 4:** Mapping data for nine consecutive reads. Reads mapped to GRCh38 using minimap2 -x map-ont. The combined read length is 1,385,925 bases and maps to a total span of 1,470,878 bases.

| Read ID | No. | Ch | Len | Chr | Std | Start | End |
| --- | --- | --- | --- | --- | --- | --- | --- |
| 3ce0651d-5a64-4d57-bd5c-3cd57045d473 | 325 | 56 | 262,945 | 13 | - | 46000683 | 46279489 |
| 6588b110-ef2c-48a2-b722-74902345f0dd | 329 | 56 | 53,061 | 13 | - | 45944001 | 46000240 |
| 5e3eeb11-9f03-45c5-b00e-99bdab5c1c3d | 333 | 56 | 21,821 | 13 | - | 45921320 | 45943803 |
| 07a93f40-2e33-451d-bb00-623d5bdebdfd | 338 | 56 | 3,096 | 13 | - | 45917140 | 45920055 |
| 1fac5b9f-9a73-45e4-a8d1-5336161f08c6 | 339 | 56 | 164,502 | 13 | - | 45742838 | 45916806 |
| c6a7d4b5-252b-4665-bc1e-77a315154edc | 340 | 56 | 300,951 | 13 | - | 45424411 | 45742140 |
| c0d044e3-668d-4794-a535-c77a16fb850a | 344 | 56 | 94,514 | 13 | - | 45323669 | 45423042 |
| 491016a6-b817-4af9-8fd5-9a3572326c95 | 348 | 56 | 370,342 | 13 | - | 44930258 | 45320732 |
| 8ed6cd0a-f4e6-4785-810d-f8193688bdf0 | 350 | 56 | 114,693 | 13 | - | 44808611 | 44929104 |

Investigating these reads in more detail revealed that changes in the normal current flow can be seen which appear to cause the real time MinKNOW read detection to split the read. Occasionally, these events trigger unblock activity, after which the read continues to sequence from the same point in the reference. We observed one instance where this unblock loop lasted in excess of 40 minutes and then continued to sequence the same molecule (Supplementary Figure 3). The most complex fused read observed to date consists of 38 individual reads which all map contiguously to the same region of the genome (Figure 3D, Supplementary Figure 4, Supplementary File Collection 2). In fact, the read illustrated in Figure 1 also represents a ‘fused read’. When called as a single read, the base called sequence maps contiguously to chromosome 1 from 60,882,202 to 61,129,414 bases (spanning 247,212 bases).

Parsing data from within a representative bulk fast5 file from this more recent data set identified a number of annotation states that appear to correlate with the starts and ends of incorrectly split reads (Figure 4). Predominantly these reads show either ‘above’ or ‘transition’ classifications occuring at the change from one read to the next. At a much lower frequency, we observed unblocks splitting reads. These ‘above’ or ‘transition’ signals can be clearly seen in the signal traces (see Figure 2). We wondered if interference from surrounding channels might cause this. However, grouping signals from the immediately surrounding channels failed to reveal any clear pattern (data not shown). We currently have no clear explanation for why these reads are incorrectly split. We note that the frequency of read splitting varies between DNA extractions.

**Figure 4.**
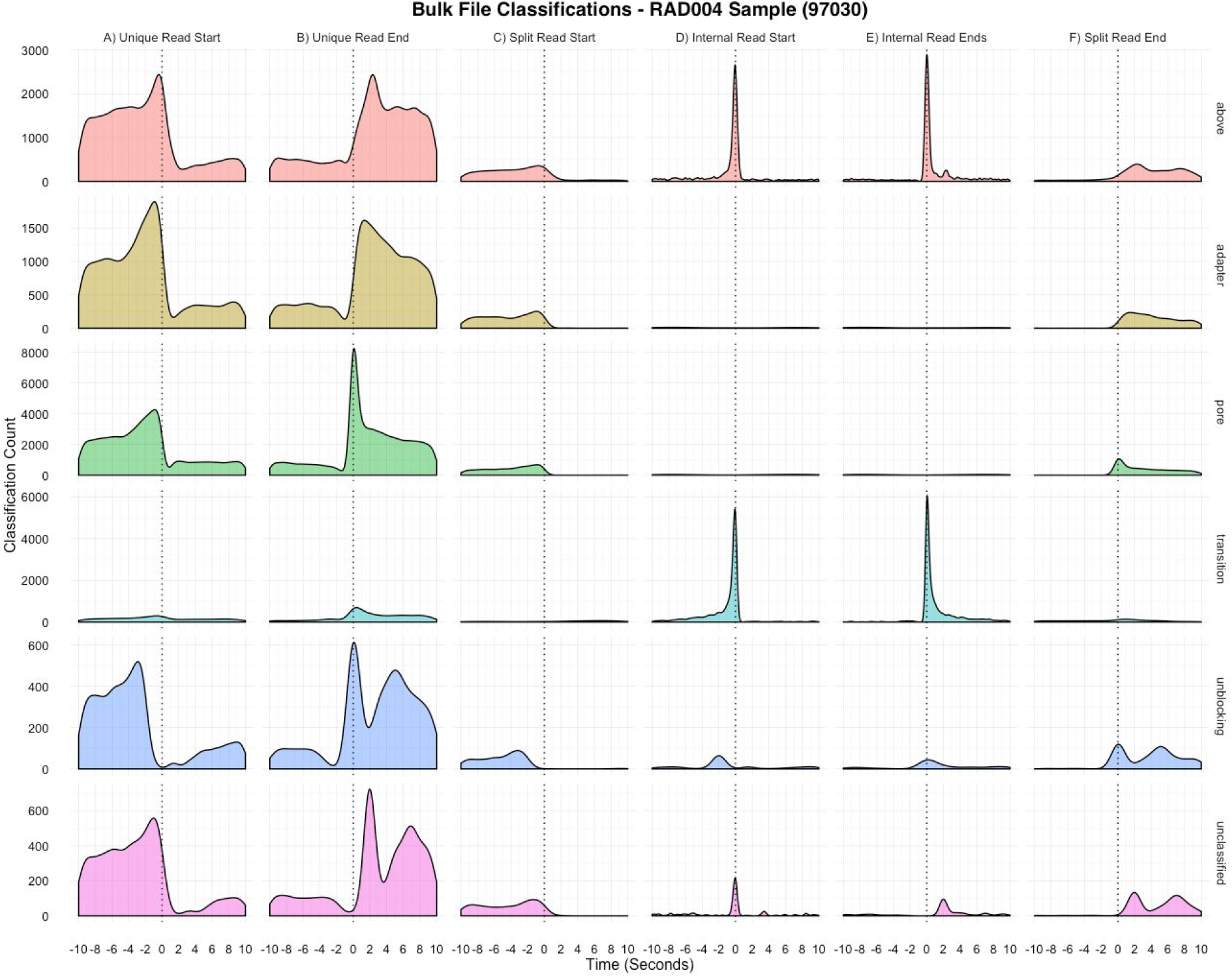
MinKNOW Classifications. A selection of classifications captured from a bulk fast5 file. Reads are identified as unique reads (i.e not fused) (A,B) and then split reads (C-F). MinKNOW classifications in the ^10^ seconds before and after a read start or end are counted for every read type. The abundance of ‘above’ and ‘transition’ classifications at the incorrect read splits by MinKNOW can be clearly seen.

## Discussion

BulkVis is a basic tool for visualising bulk fast5 files collected from Nanopore sequencers. As a consequence of developing BulkVis, we have identified that ultra-long reads can be incorrectly split by MinKNOW resulting in artificially shorter reads. This appears to disproportionately affect ultra long read preparations, although this requires further investigation. We note that the method we use for generating ultra long reads is outside the range of normal operating conditions for nanopore sequencing as recommended by Oxford Nanopore ^11^. Also, the number of ultra long datasets analysed in this way is limited at this time. However, for those wishing to maximise read length the fact that adjacent reads from a single pore may represent a single molecule of DNA is likely to be of great interest. Although we have no formal explanation for why this read splitting occurs, we speculate that in some cases this might be caused by DNA damage or contaminants physically linked to the DNA causing spikes in the signal seen by MinKNOW. We cannot exclude the possibility that some of the split reads we observe are caused by single strand breaks with the DNA after the break being captured by the pore.

An additional observation is that in some cases reversal of the voltage does not successfully reject a read. We emphasise that this effect is apparently rare and typically occurs within long reads. For applications such as selective sequencing ^5^, reads will be rejected early in the sequencing process. We argue that this will be far more efficient than reads that are rejected midway through their length or are rejected due to some linked contaminant blocking the pore. This aligns with our previous observations on ‘read until’ ^5^. However, it is impossible to quantify the precise nature of reads that are successfully rejected.

We provide helper scripts to identify candidate incorrectly split reads. However, these scripts are limited as they currently rely on a suitable reference genome to map reads against. We believe it is possible to recognise candidate reads by close analysis of bulk fast5 files, but in the future we suspect that MinKNOW itself can be further optimised to avoid these incorrectly split reads. There is a tension between under splitting reads, which could lead to chimeras ^13^ and over splitting which results in the artificially shortened reads seen here. For general use, over splitting is clearly preferential to incorrect chimeras. However, those interested in assembly and maximising long reads should be aware that dynamic decisions made by MinKNOW in splitting reads may not always be correct. It might in future be possible to identify candidate incorrectly split reads from the absence of adapter sequences at the start of a read.

Whilst we see no requirement for the routine collection of bulk fast5 files from Nanopore sequencers, those interested in de novo assembly of genomes in the absence of a reference may benefit from collecting these files. BulkVis is provided for the visual inspection of challenging or difficult to sequence samples or where the user wishes to investigate specific events during a run. In these instances analysis of a bulk fast5 file may provide some visual indication of the underlying issues.

## Methods

### Sequencing

Sequencing was carried out using high molecular weight DNA extracted and prepared for sequencing as described by Josh Quick ^8,11^. RAD002 datasets are as described in ^8^. RAD004 sequencing was performed using MinKNOW version 1.11.5. Standard MinKNOW running scripts were used with regular manual restarting to maximise the number of sequencing channels. Restarts were timed to every two hours, to coincide with the 5mV change in voltage during sequencing. Thus the theoretical maximum read length is 3.24 Mb.

### Installation

BulkVis and companion scripts are available on github (at https://www.github.com/LooseLab/bulkvis). The scripts make use of the python modules: NumPy ^14^ , Pandas ^15^, bokeh ^16^ and h5py ^17^. Full instructions and documentation are provided at http://bulkvis.readthedocs.io.

### Operation

BulkVis is started from the command line, with the current working directory as the parent of bulkvis, using `bokeh serve --show buikvis`. This will start the server and load the application in the user’s default web browser.

### Companion Scripts

#### Prerequisites

Mapping files for the following scripts are generated using minimap2. Scripts expect the default paf format output from minimap2 using the -x map-ont option ^18^. Some scripts require a bulk fast5 file which must be collected according to instructions from Oxford Nanopore Technologies available from the Nanopore Community ^6^. The scripts require a sequencing_summary.txt file as generated by albacore or guppy. Most script dependencies are installed with BulkVis, scripts that require other tools are stated below.

`minimap2 -t 4 -x map-ont /path/to/reference.fa */*.fastq > mapping.paf`

##### 1. whale_watch.py

This script takes as input a single sequencing_summmary.txt file generated by albacore or guppy (Oxford Nanopore base callers) and a paf format output file from minimap2. Optionally the user can select the distance threshold between read ends and starts for them to be considered a single molecule which defaults to 10 kb. The script outputs statistics on the input reads, the means, medians and N50s with and without correction for read splitting and a list of the top read lengths seen in an individual run. The script writes a file (default fused_reads.txt) that lists every fused read in the dataset. It provides the coordinates of the fused read in a format compatible with BulkVis to enable quick viewing.

`python3 whale_watch.py -s /path/to/sequencing_summary.txt -p /path/to/mapping.paf`

##### 2. whale_merge.py

This script builds on whale_watch.py to generate a fastq file for each detected fused read in a sequencing_summary.txt file. It requires a folder of fastq files in addition to the sequencing_summary.txt file and a relevant .paf file. It can either output all the reads from a run including corrected fused reads and all non-fused reads, or just output the corrected fused reads.

`python3 whale_merge.py -s /path/to/sequencing_summary.txt -p /path/to/mapping.paf -f /path/to/fastq/reads/`

##### 3. whale_plot.py

This script is built on whale_watch.py and requires a sequencing_summary.txt, .paf file, and bulk fast5 file to produce six CSV files containing the distributions of MinKNOW events around read starts and ends. Whale_plot.py then calls whale.R to produce a plot similar to Figure 4. Rscript and packages ggplot2, dplyr, and tidyr are required to produce the plot.

`python3 whale_plot.py -s /path/to/sequencing summary.txt -p /path/to/mapping.paf -b /path/to/bulkfile.fast5 -t 10`

##### 4. pod_plot.py

This script generates plots for all reads in a fused_reads.txt file. This uses bokeh to render a plot (in a headless browser) and requires selenium and Pillow to be installed from pip/conda; and phantomjs from http://phantomis.org/.

`python3 whale_watch.py -f /path/to/fused_reads.txt -b /path/to/bulkfile.fast5 -D output_folder`

##### 5. bulk_info.py

Given a directory containing bulk fast5 files this script outputs a CSV containing run information. Attributes reported are: sample frequency, run id, experiment, flowcell id, protocol version, minknow version, minion id, hostname, sequencing kit, flowcell type, asic id and experiment start time.

`python3 bulk_info.py -d /path/to/folder/of/bulkfiles/`

## Acknowledgements

This work was supported by BBSRC grant (BB/N017099/1 and BB/M020061/1), the Wellcome Trust (204843/Z/16/Z) and a BBSRC iCASE studentship (BB/M008770/1,1949454). The authors thank members of DeepSeq at Nottingham, Nick Loman, Josh Quick, Jared Simpson and John Tyson for helpful discussions and advice as well as Graham Hall (Oxford Nanopore Technologies) for insights into unblocking behaviour.

## Supplementary Materials

**Supplementary Note 1**

The bulk fast5 file used here can be downloaded from http://s3.amazonaws.com/nanopore-human-wgs/bulkfile/PLSP57501_20170308_FNFAF14035_MN16458_sequencing_run_NOTT_Hum_wh1rs2_60428.fast5

**Supplementary Table 1.**
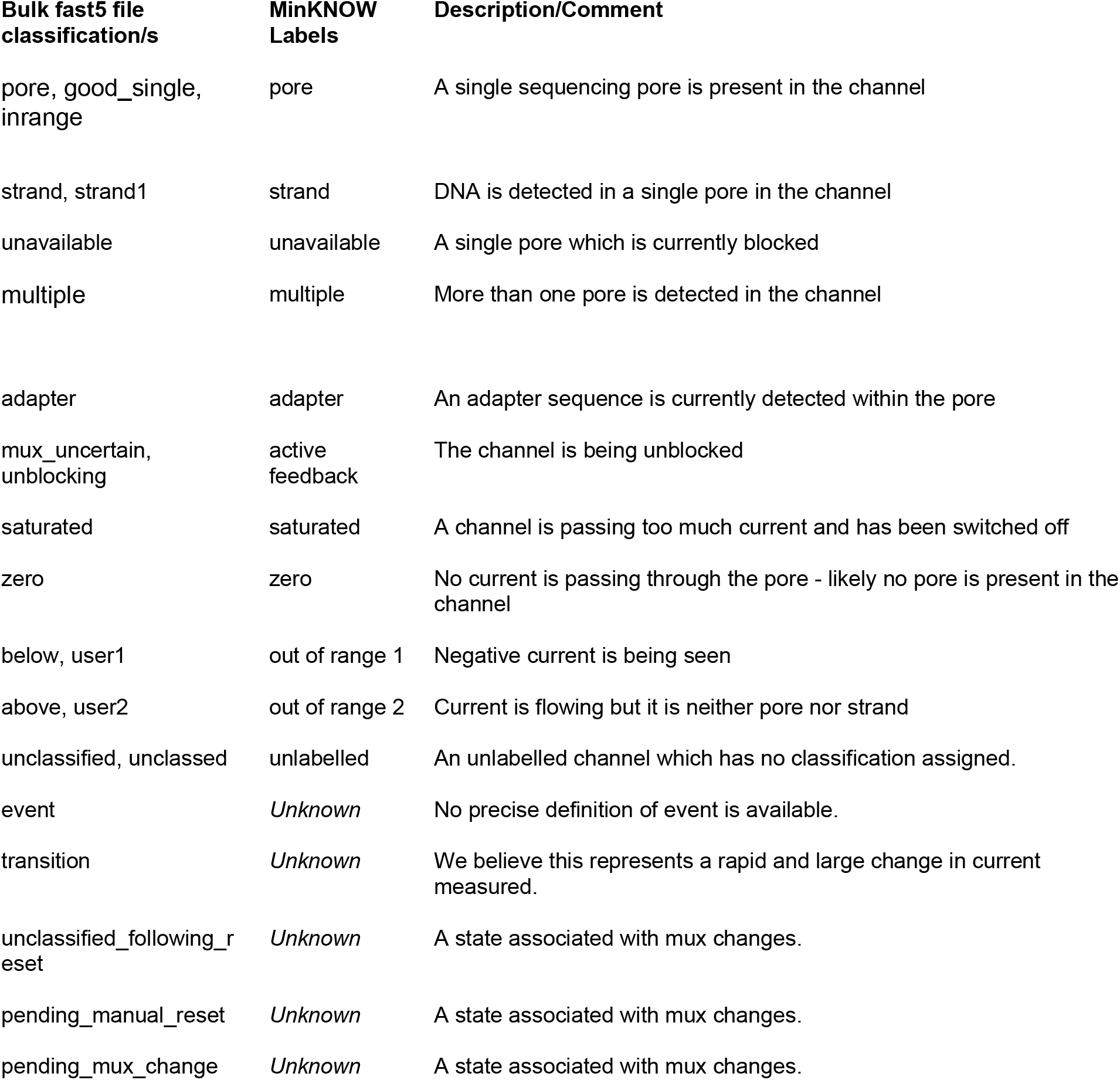
Classification Descriptions. MinKNOW labels described on the Oxford Nanopore Forum (https://community.nanoporetech.com/support/faq/test1/minknow/minknow/what-are-the-colours-shown). To our knowledge there is no detailed description of the relationship between bulk fast5 file classifications and MinKNOW labels. This table presents our assumptions about the relationship between bulk fast5 labels and MinKNOW classifications.

**Supplementary Figure 1.**
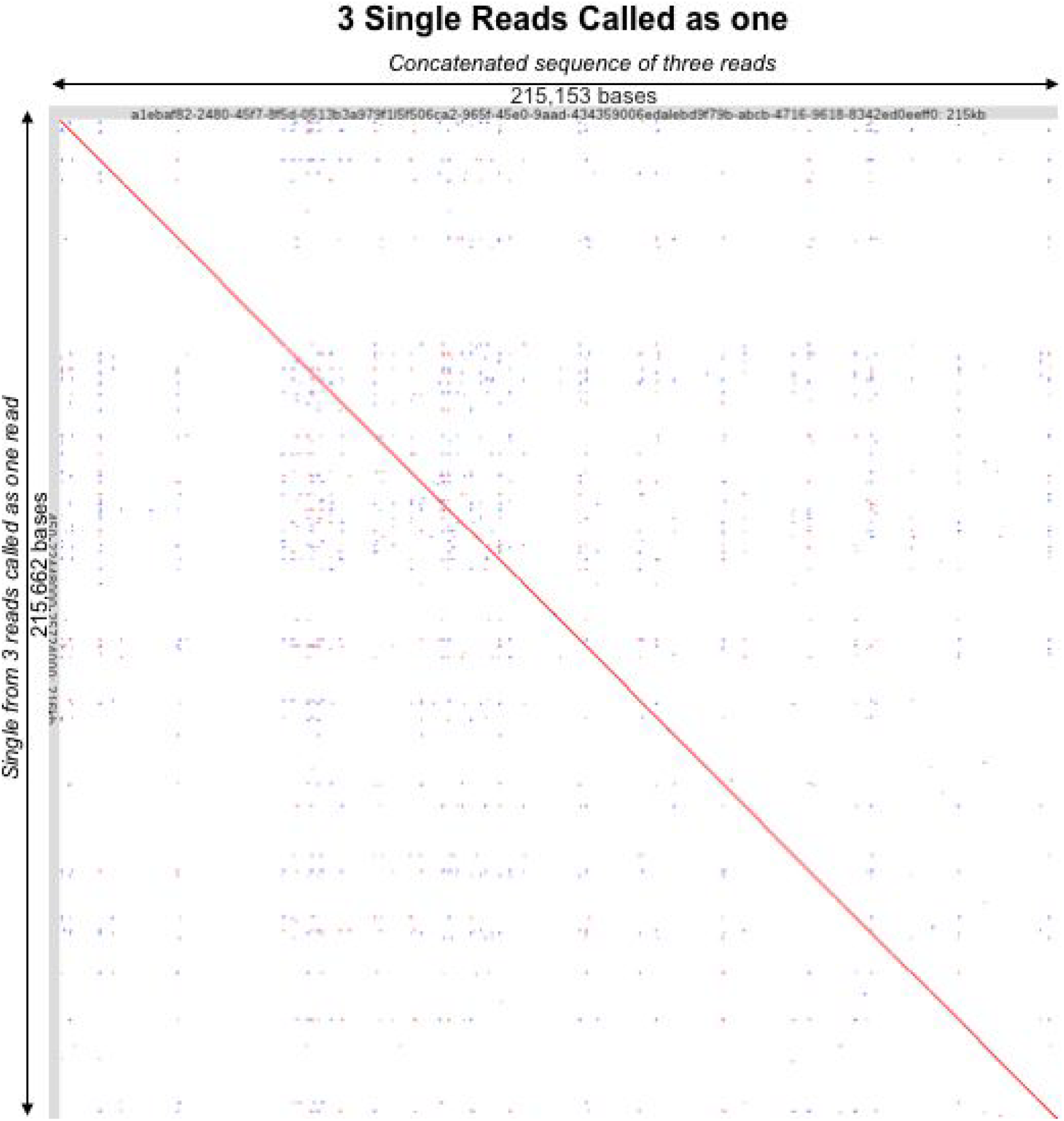
A simple last alignment and dot plot of the three individual basecalled reads shown in Figure 1 aligned against the merged signal for each of those three reads called as one read by BulkVis.

**Supplementary Figure 2.**
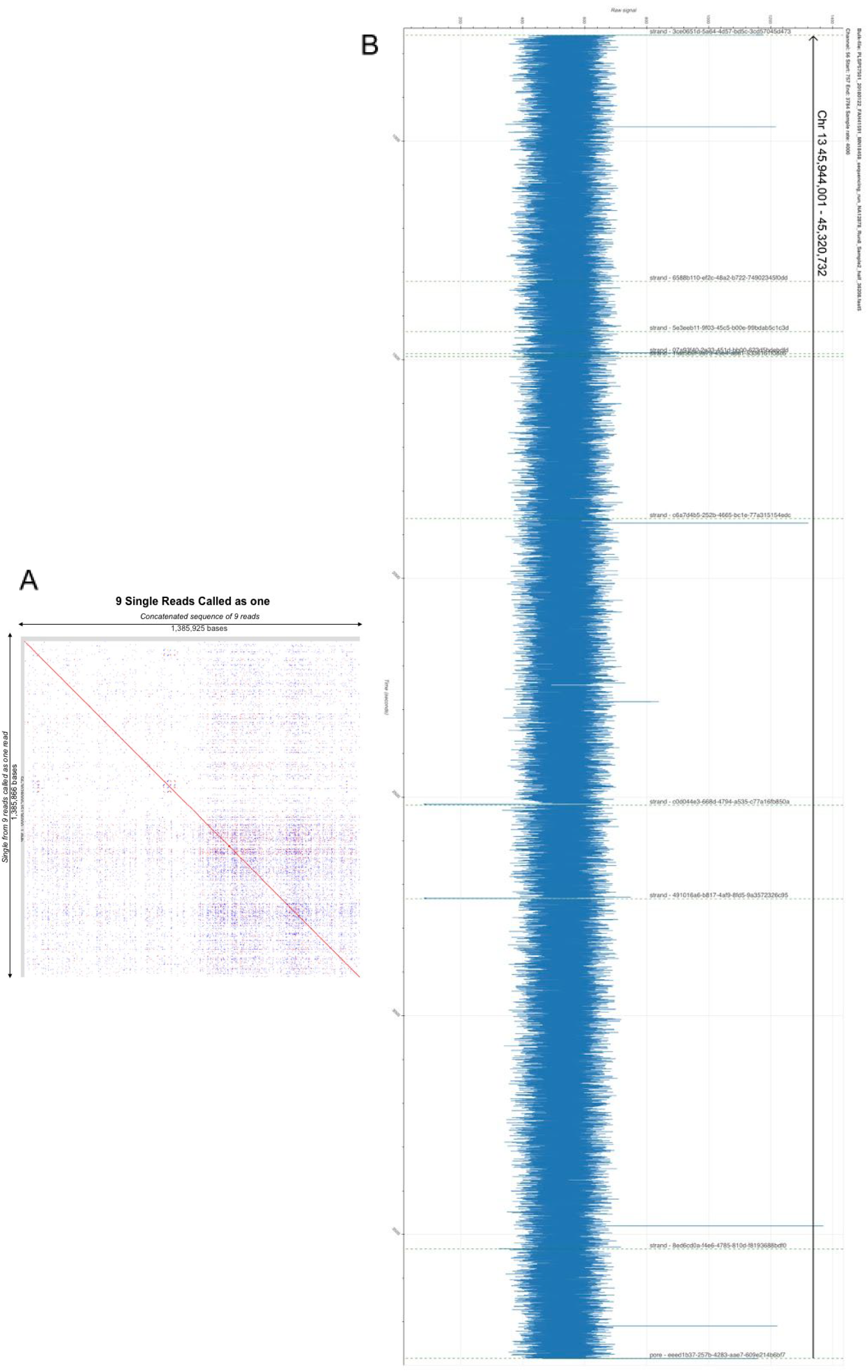
A) Mapping the concatenated basecalled reads against the single read called from squiggle by BulkVis. B) The full length signal region spanning 9 individual reads from a bulk fast5 file as shown in Figure 3C. Dashed lines indicate new reads identified by MinKNOW.

**Supplementary Figure 3.**
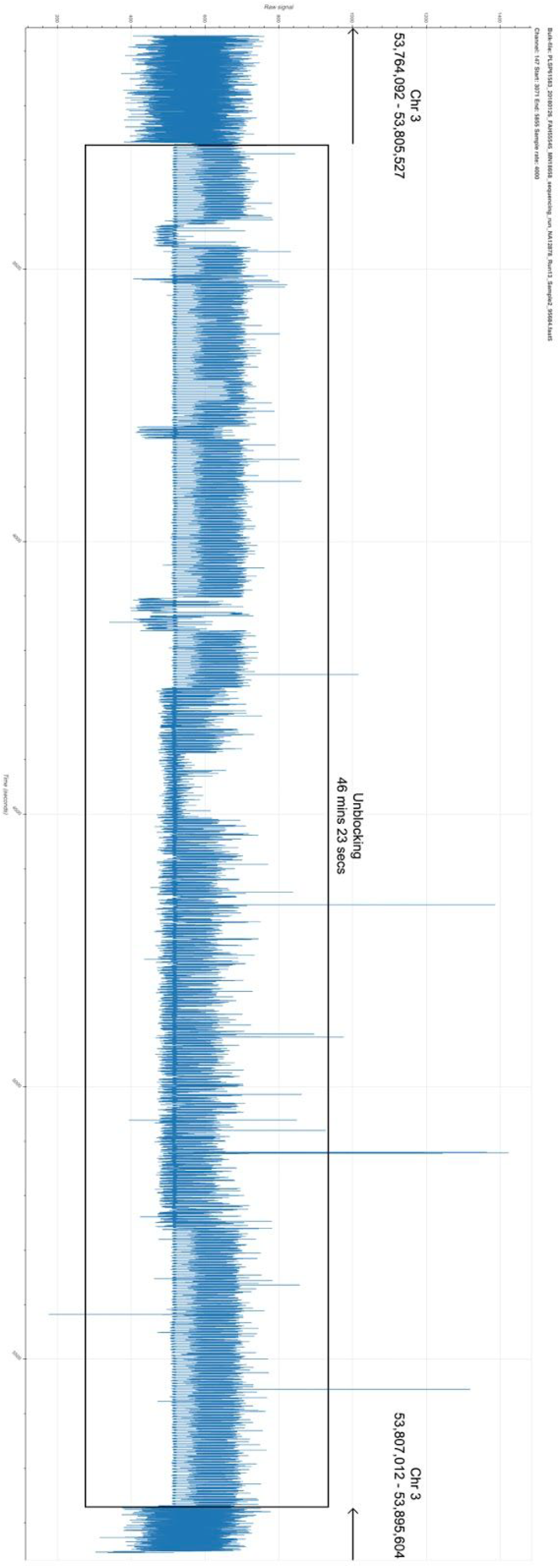
A read containing an unblock sequence which lasts over 46 minutes, but later continues to sequence the same molecule. Dashed lines indicate new reads identified by MinKNOW. The boxed region contains repeated unblocks.

**Supplementary Figure 4.**
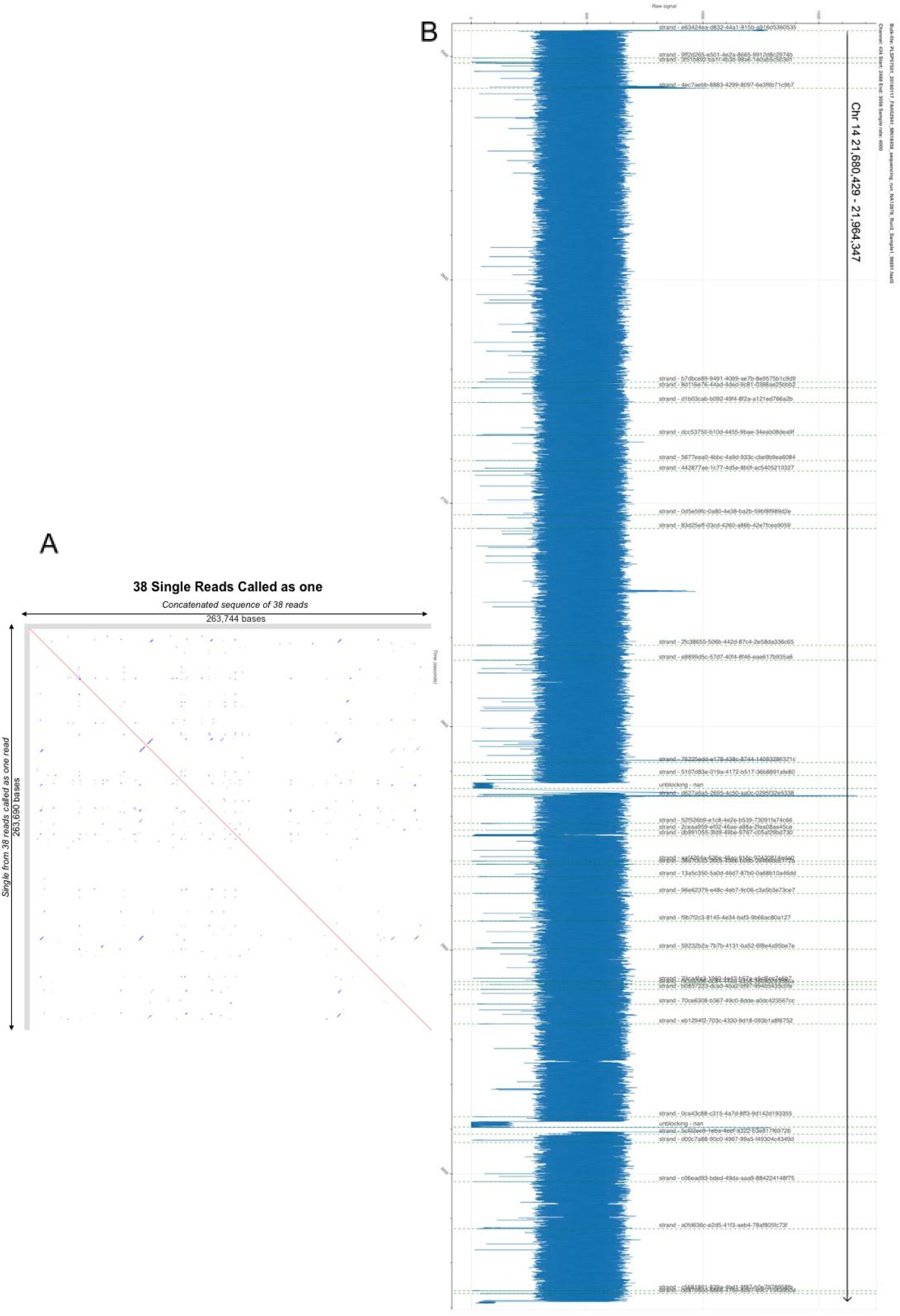
A) Mapping the concatenated basecalled reads against the single read called from squiggle by BulkVis. B) The full length signal region spanning 38 individual reads from a bulk fast5 file as shown in Figure 3D. Dashed lines indicate new reads identified by MinKNOW.

**Supplementary File Collection1**

This file is a tar.gz archive containing the sequencing_summary.txt and mapping.paf from the MinION with ASIC ID 3976726082. The ‘bulkvis’ folder contains a read fast5 file generated by BulkVis and two fastq files that were created by calling the BulkVis read fast5 file and merging the original split fastq files; the ‘original’ folder contains the read fast5 files as originally split by MinKNOW and their resulting fastq file.

**Supplementary File Collection2**

This file is a tar.gz archive containing four subfolders: 3A_longest-single-read, 3B_longest-fused-read-without-bulkfile, 3C_longest-fused-read-with-bulkfile, and 3D_read-with-38-splits; each of these folders holds an ‘original’, ‘bulkvis’, or both folders. The ‘original’ folders contains the read fast5 files as originally split by MinKNOW and their resulting fastq files; the ‘bulkvis’ folders contains read fast5 files generated by BulkVis and fastq files that were created by calling the BulkVis read fast5 file and merging the original split fastq files.

